# Wired together, change together: Spike timing modifies transmission in converging assemblies

**DOI:** 10.1101/2023.05.22.541797

**Authors:** Lidor Spivak, Shirly Someck, Amir Levi, Shir Sivroni, Eran Stark

## Abstract

Precise timing of neuronal spikes may lead to changes in synaptic connectivity and is thought to be crucial for learning and memory. However, the effect of spike timing on neuronal connectivity in the intact brain remains unknown. Using closed-loop optogenetic stimulation in CA1 of freely-moving mice, we generated new spike patterns between presynaptic pyramidal cells (PYRs) and postsynaptic parvalbumin-immunoreactive (PV) cells. This stimulation led to spike transmission changes which occurred together across all presynaptic PYRs connected to the same postsynaptic PV cell. The precise timing of all presynaptic and postsynaptic cells spikes impacted transmission changes. These findings reveal an unexpected plasticity mechanism, wherein spike timing of a whole cell assembly has a more substantial impact on effective connectivity than that of individual cell pairs.

## Introduction

At the core of our capability for learning and memory is the capacity of the brain to adapt and modify in accordance to external events^1, 2^. Learning is supported by changes in synaptic connections between neurons, modulated by different plasticity rules^3, 4^. One model, spike timing-dependent plasticity (STDP), posits that changes in synaptic connectivity are driven by the relative timing of spikes between pre- and postsynaptic neurons^5–8^. In vitro experiments showed that the millisecond-timescale of spike timing between a pair of neurons influences their synaptic connectivity^9–12^. However, the experiments did not reveal whether similar plasticity rules apply in the intact brain, where numerous cells are active simultaneously.

STDP studies in intact animals typically involved pairing the activity of a single postsynaptic cell with either sensory^13–15^ or optogenetic stimuli^16, 17^. However, external stimuli activate an entire presynaptic pool rather than a single presynaptic neuron as done in vitro. Additionally, these studies assessed plasticity changes based on the response of the postsynaptic cell to the external stimuli, neglecting spike timing and alterations of individual connections between the cells. Thus, the impact of spike timing of individual pairs within the same assembly on their connectivity remains unclear.

## Results

### Changes in spike transmission gain occur after closed-loop induction of PV spikes

To investigate how spike timing affects connectivity in the intact brain when multiple neurons are involved, we recorded the simultaneous activity of dozens of PYRs and PV interneurons in mouse CA1. The PV cells receive excitatory input from multiple PYRs, demonstrating a wide range of connection strengths^18–21^. Together, this set of presynaptic PYRs and their postsynaptic target PV cell form a converging assembly (CA; Fig. 1A). The CA architecture makes the PYR-to-PV interface useful for testing how spike timing changes neuronal connectivity with respect to other connections.

**Figure. 1.**
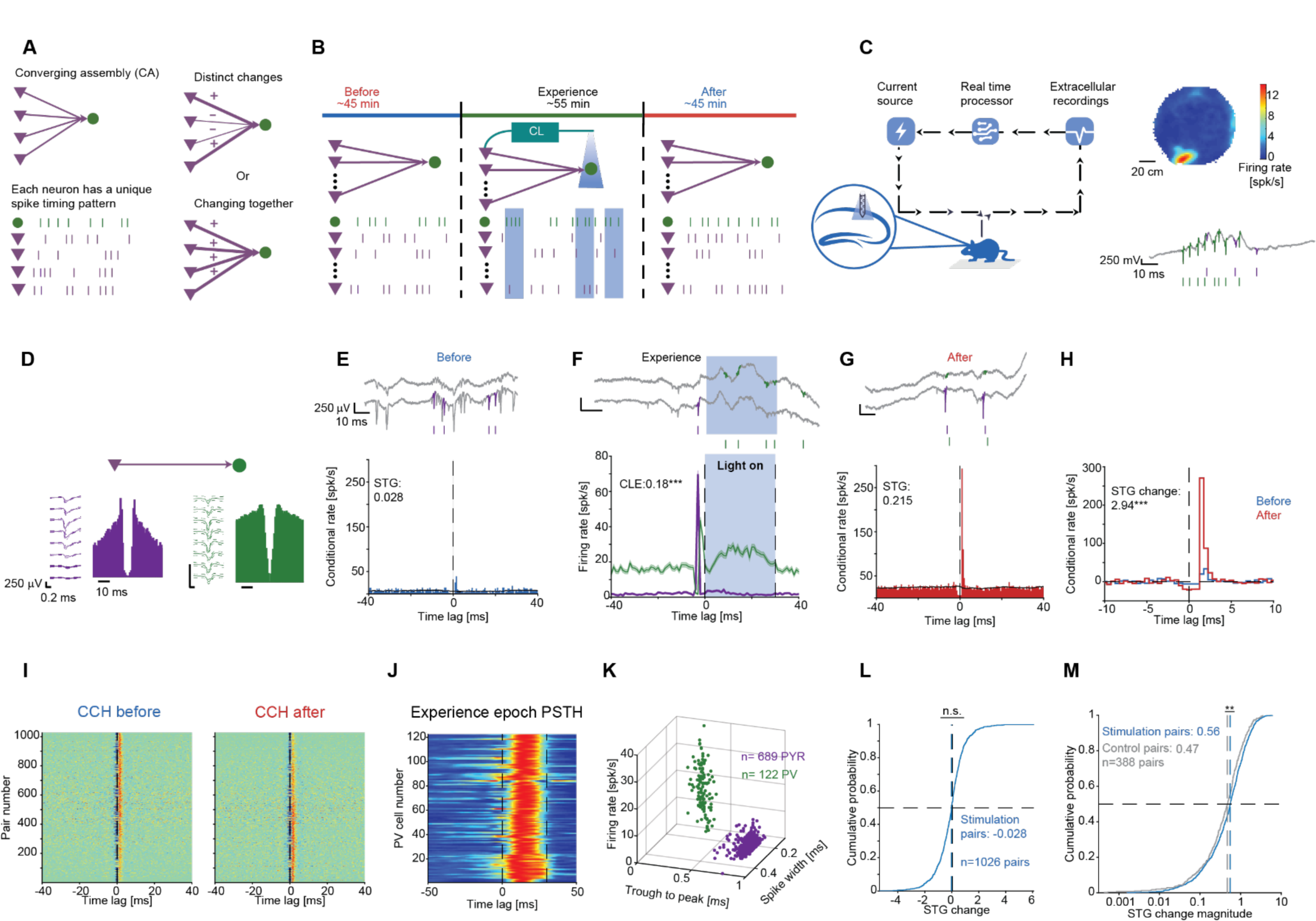
Changes in spike transmission gain occur after closed-loop induction of PV spikes. (**A**) The STG between two neurons in a CA may be affected by their spike timing, spike timing of the whole CA, or be spike timing history independent. (**B**) Experimental paradigm. (**C**) Closed-loop system. **Left**, During the “Experience” epoch, spikes of one or more PYRs detected in real-time generate light-induced spiking in nearby PV cells. **Right**, Place field of a PYR on the open field, and wideband (0.1-7,500 Hz) trace during a ripple event. (**D**) Wideband waveforms and autocorrelation histograms (ACHs) of a pre-postsynaptic PYR-PV pair. (**E**) Conditional rate cross-correlation histogram (CCH) during “Before”. **Top**, Wideband traces. (**F**) Experience epoch peristimulus time histograms (PSTHs). PV firing rate increases immediately after PYR spiking, and again during the light. CLE, closed-loop efficiency. (**G**) “After” epoch CCHs. (**H**) Overlaid CCHs for the no-light epochs. ***: p<0.001, permutation test. (**I**) Features of PYRs and PV cells recorded during stimulation experiments. (**J**) PSTHs of the PV cells. (**K**) All pairwise CCHs, exhibiting excitatory connectivity. (**L**) STG changes for the Stimulation PYR-PV pairs. n.s.: p>0.05, Wilcoxon’s test. (**M**) Absolute STG changes for Stimulation and Control pairs. **: p<0.01, U-test.

To determine how spike timing between multiple presynaptic PYRs and a postsynaptic PV cell influences the effective connectivity, we designed the following experiment (Fig. 1B). We recorded baseline CA1 network activity for a median of 45 min (“Before” epoch), followed by 55 min of “Experience” and an additional 45 min of baseline (“After”). During all three epochs, mice were free to behave in a familiar environment (Table S1; Fig. 1C). We manipulated PYR-PV spike timing during the “Experience” epoch, and measured the effects of the short-term spike timing changes by comparing effective connectivity changes between the “Before” and “After” epochs. To manipulate spike timing during the “Experience” epoch only, the spiking activity of one or more PYRs was detected in real time (Fig. 1D,F). After a 3 ms processing delay, a 30 ms light stimulus was given, inducing spiking in nearby PV cells. To quantify changes in effective connectivity between the “Before” and “After” epochs, we used the spike transmission gain (STG^22^) metric (Fig. 1E-G). To assess long-term changes in STG, we defined the “STG change” as the base-2 logarithm of the ratio between the STGAfter and the STGBefore (Fig. 1H), where a change of 1 or −1 signifies STG doubling or halving following the “Experience” epoch.

To understand how changes in spike timing during the “Experience” epoch affect changes in spike transmission, we focused on PYR-PV pairs which exhibit monosynaptic connectivity (p<0.001, Poisson test; Fig. 1I), and in which the PV cell was activated by the closed-loop stimulation (p<0.05, Poisson test; Fig. 1J). PYRs and PV cells could be accurately differentiated based on waveform features or spike timing statistics (Fig. 1K). A set of 689 connectivity-tagged PYRs and 122 optically-tagged PV cells yielded a cohort of 1026 Stimulation pairs, recorded during a total of 29 sessions from four freely-moving PV::ChR2 mice (Table S2). Among these pairs the median STG change was not consistently different from zero (−0.028; p=0.09; Wilcoxon’s test; Fig. 1L). Thus, there is an equilibrium of changes in STG at the population level.

To determine whether the observed changes exceed spontaneous changes, we compared the 1026 Stimulation pairs with Control pairs, recorded during long no-stimulus periods from the CA1 of five mice (Table S3). All 388 Control pairs exhibited monosynaptic connections (p<0.001, Poisson test) but were not exposed to any light stimuli. The median STG change of the Control pairs (0.004) was not consistently different from the Stimulation pairs (p=0.37, U-test). However, the magnitude of the STG changes among the Stimulation pairs (median [interquartile range, IQR]: 0.556 [0.240 1.079]) was higher than Control pairs (0.473 [0.205 0.863]; p=0.006, U-test; Fig. 1M). Thus, while the overall net STG change remains balanced, the magnitude of STG changes increases following closed-loop stimulation during the “Experience” epoch.

### Changes in spike transmission occur together in a converging assembly

To investigate STG equilibrium, we considered two scenarios. First, equilibrium is maintained by each CA, implying that net STG changes for every assembly are near-zero. Second, equilibrium is maintained only at a higher level, where some CAs exhibit a net STG increase and others a decrease. To distinguish between the scenarios, we compared the STG of individual pairs to other pairs within the same assembly, referred to as “peers” (Fig. 2A). To control for more global changes including influences of behavior or brain state changes on STGs, we compared the STG of the individual pair to every PYR-PV pair within other simultaneously recorded CAs, referred to as “non-peer pairs” (Fig. 2A).

**Figure 2.**
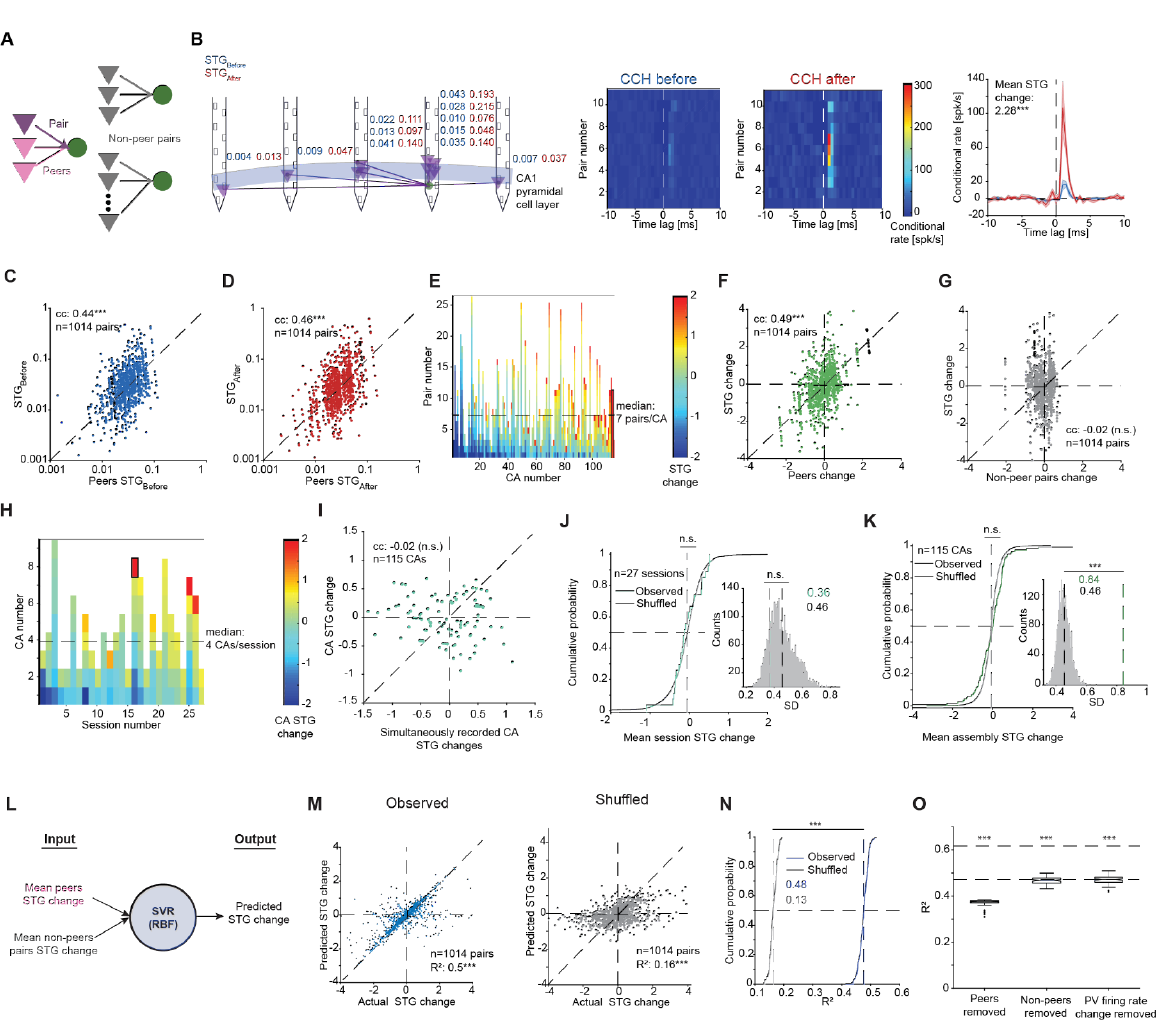
Changes in spike transmission occur together in a converging assembly. (**A**) Peer/non-peer nomenclature for simultaneous-recorded PYR-PV pairs based on their relation to a specific pair. (**B**) Example 12-unit CA, CCHs for the 11 pairs, and Average CCHs. Here and in **O**, ***: p<0.001, Wilcoxon’s test. (**C**) Pairwise STGBefore vs. the mean peer STGBefore. Here and in all other panels, n.s./***: p>0.05/p<0.001, permutation test. (**D**) Same, for STGAfter. (**E**) Pairwise STG changes per CA. (**F**) Pairwise STG change vs. mean STG change of all peers. (**G**) Same, vs. mean non-peer STG change. (**H**) CA STG changes per session. (**I**) CA STG change vs. mean STG change of all other simultaneously-recorded CAs. (**J**) Mean STG changes of same-session pairs and chance distribution. **Right**, SD of same-session STG changes. (**K**) Mean STG changes of all multi-pair CAs and chance distribution. **Right**, SD of CA STG changes. (**L**) Cross-validated SVR. (**M**) **Left**, Predicted vs. actual STG changes. **Right**, Same, for intra-session shuffled allocation. (**N**) R^2^ values derived from 100 independently-generated SVRs. (**O**) Contribution of postsynaptic firing rate to STG change prediction. Every box plot shows the median and IQR, whiskers extend for ±1.5 IQRs, and individual dots indicate outliers.

We found that during both the Before and the After epochs, single-pair STG was correlated with the mean peers STG (Before: rank correlation coefficient [cc]: 0.44; p<0.001, permutation test; After: cc: 0.46; p<0.001; Fig. 2C-D). The correlation between single-pair STG and peers STG was also observed in two other datasets, recorded from CA1^22^ (cc: 0.75; p<0.001; Fig. S1A) and neocortex^20^ (cc: 0.46; p<0.001; Fig. S1B). In contrast, single-pair STG did not consistently correlate with the mean STG of non-peer pairs in neither the "Before" (cc: 0.01; p=0.902) nor the "After" epoch (cc: −0.05; p=0.080; Fig. S1C-D), indicating that CA STG similarities extend beyond brain state changes. Furthermore, PYR-to-PYR firing synchrony was higher for same-CA pairs compared with different-CA pairs (p<0.001, Wilcoxon’s paired test; Fig. S1I-J). Consequently, CA PYRs exhibits synchronous firing and similar STGs with the same postsynaptic PV cell.

To determine whether STG change of a specific PYR-PV pair correlates with other same-CA changes, we compared for each pair the mean STG change of same-assembly peers (Fig. 2E). STG change of a single pair was correlated with the peer pairs STG change (cc: 0.49; p<0.001, permutation test; Fig. 2F) but not with the non-peer STGs (cc: −0.02; p=0.59; Fig. 2G). Similar results were observed in Control pairs which did not receive any light stimulation (Fig. S1E-H). Thus, PYR-PV pairs belonging to the same CA exhibit similar STG changes.

To determine whether the mean STG of individual CAs is balanced at the session level, we compared the mean STG change of each CA to the mean STG of simultaneously recorded CAs (Fig. 2H). The CA STG change did not consistently correlate with the mean STG change of other CAs (cc: −0.02; p=0.58, permutation test; Fig. 2I). Moreover, STG changes of simultaneously recorded CAs were balanced, and similar to sham CAs constructed by randomly shuffling the allocation of STGs to CAs (mean: −0.06, chance mean: −0.06, p=0.53; SD: 0.36, chance SD: 0.46, p=0.863; n=27 sessions; Fig. 2J). However, inter-assembly variability of CAs (0.84) was higher than random CAs (SD, 0.46; p<0.001, permutation test; Fig. 2K), indicating that STGs of same-CA pairs increase or decrease together. Thus, STGs of PYR-PV pairs that belong to the same CA change in a coordinated manner while preserving equilibrium of the STG changes over multiple assemblies.

To quantify the predictive power of the assembly structure to STG changes we employed cross-validated support vector regression (SVR; Fig. 2L). In one SVR, we used the mean STG change of the peers and non-peers as an input (Fig. 2M, **left**). For another SVR, we used randomly shuffled pairs from the same session (Fig. 2M, **right**). The reconstruction-based R^2^ of the individual PYR-PV STG change yielded by the first SVR was 0.48, consistently higher than the R^2^ yielded by the shuffled SVR (0.13; p<0.001, U-test; Fig. 2N).

Finally, we examined whether the predictive power of the assembly structure extends beyond postsynaptic firing rate changes. Alterations in the postsynaptic cell firing rate from the Before to the After epoch were consistently correlated with STG changes (cc: 0.535; p<0.001; permutation test; Fig. S1K). To evaluate the individual contributions of peer pairs, non-peer pairs, and postsynaptic firing rate changes to predicting STG changes of a specific PYR-PV pair, we trained three separate SVRs while removing one feature at a time. R^2^s produced by all three SVRs were lower compared with the full model (trained using all three features; p<0.001, Wilcoxon’s paired test; Fig. 2O), and the lowest R^2^ was observed when peer pairs information was removed. Thus, STG changes of a single PYR-PV pair can be predicted by considering the assembly structure.

### Spike timing of the whole converging assembly predicts changes of spike transmission gain

To determine whether spike timing influences the changes in STG within CAs, we examined the impact of immediate spike timing alterations during light stimulation on STG changes. During stimulation, PV cells firing rates increased, exhibiting a light-induced firing rate gain of 1.56 [1.29 2.55] (n=122; Fig. S2A). 198/689 (29%) PYRs were “trigger” PYRs, exhibiting a consistent PSTH peak 3 ms [3 3] ms before light onset (Fig. 3A), with a closed-loop efficiency (CLE) of 0.054 [0.013 0.184] (Fig. S2B-C). However, consistent PSTH peaks were not associated with higher STG changes (Fig. 3B), and the CLE of the PYRs did not consistently correlate with individual STG changes (cc: 0.02; p=0.42; permutation test; Fig. 3C). Furthermore, PV cell firing rate gain did not correlate with the mean CA STG changes (cc: −0.10; p=0.69; Fig. 3D). Thus, the responses of the individual cells to light stimulation do not demonstrate a consistent correlation with the STG changes.

**Figure 3.**
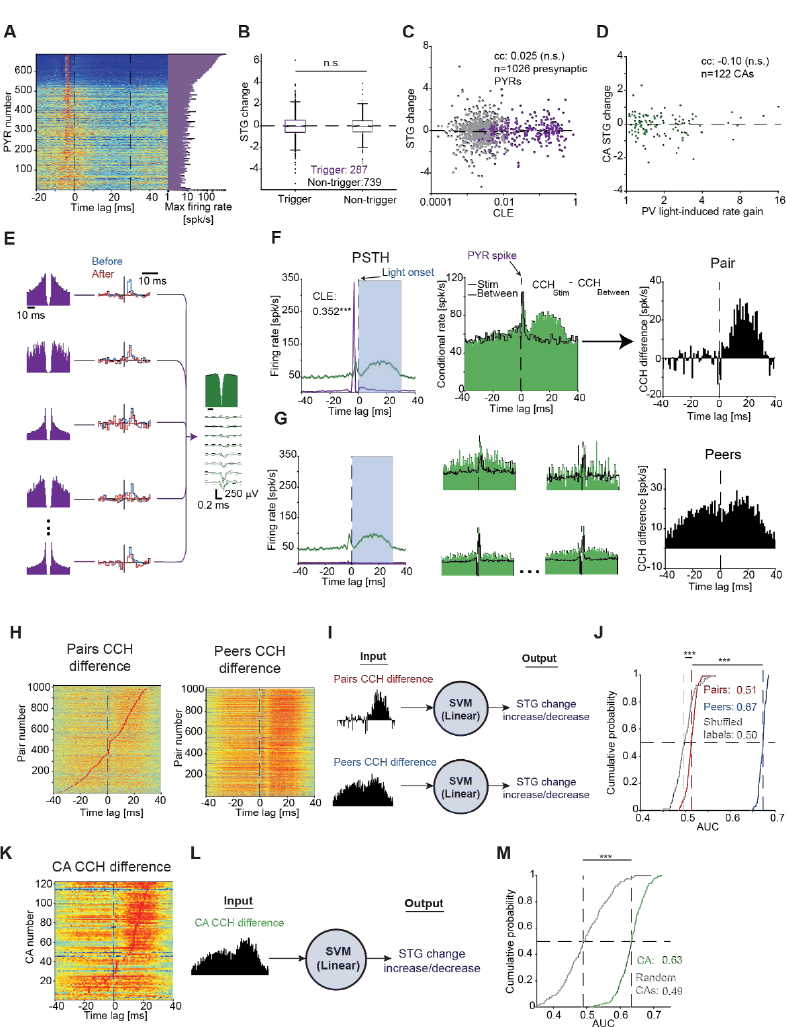
**Spike timing of the whole converging assembly predicts changes of spike transmission gain**. (**A**) PSTHs of all PYRs. (**B**) STG changes grouped by the presynaptic PYR (trigger/non-trigger). n.s.: p>0.05, U-test. (**C**) STG changes vs. CLE. Here and in **D**, n.s.: p>0.05, permutation test. (**D**) Mean CA STG change vs. PV light-induced gain. (**E**) Example CA with 11 PYR-PV pairs, five shown. (**F**) Quantification of short-term spike timing changes. **Left**, PSTHs of the top PYR-PV pair, in which the PYR triggered closed-loop PV illumination. ***: p<0.001, Poisson test. **Center**, CCHs for the PYR-PV pair during illumination (“Stim”) and in the lack thereof (“Between”). **Right**, The CCH difference quantifies the effect on the two spike trains. (**G**) **Left**, PSTHs of four other PYRs and the PV cell. **Center**, CCHs. **Right**, Summing all peer pairwise CCH differences yields the peers CCH difference. (**H**) Pairwise and peer CCH differences. (**I**) Cross-validated binary classifier trained to predict increase/decrease of pairwise STG. (**J**) AUC of 100 independently-generated classifiers. Here and in **M**, ***: p<0.001, U-test. (**K**) CA CCH differences. (**L**) CA classification. (**M**) AUC distributions for the CA classifiers.

We explored whether the interaction of PYRs and PV cell spike timing during light stimulation could predict STGs changes. In a representative CA, 11 PYRs converged on the same PV cell (Fig. 3E). Each pair may have experienced different spike patterns, affecting STG changes in distinct manners. Due to the correlation of STG changes within a CA (Fig. 2), we investigated how single-pair STG is affected by spike timing of itself and the peers. To quantify the light-induced changes in spike patterns of one pair, we computed the CCH of a single PYR-PV pair during light stimuli times (Fig. 3F). To quantify ongoing baseline patterns, we computed the CCH for the same pair using spikes that occur between stimuli (without illumination) during the Experience epoch. The difference between the CCHs during and between stimuli yields the “CCH difference” (Fig. 3F), capturing the specific effect of stimuli on spike timing during the Experience epoch. To quantify the effect of the stimuli on spike timing of the peers, we repeated the process for all other pairs in the same CA and summed all individual CCH differences (Fig. 3G). For each pair participating in a CA with at least two presynaptic PYRs, we computed both the pair CCH difference and the peer CCH difference (1020 PYR-PV pairs; Fig. 3H). Consequently, the changes in spike timing resulting from light stimulation can be captured for a single pair and its peers using the CCH difference.

To quantify the effect of light-induced spike timing changes on the single-pair STG changes, we trained cross-validated classifiers (support vector machines, SVM) to predict whether single-pair STG change increases or decreases (Fig. 3I). The first classifier used the single pair spike timing changes (pair-wise CCH differences; n=1020), and the second classifier used the peer pairs spike timing changes (peers CCH differences). The classifier that used peer pairs CCH differences yielded higher AUCs, 0.673 [0.664 0.677], compared with the pairs classifier (AUC: 0.513 [0.506 0.521]; p<0.001, U-test; Fig. 3J). Similar results were observed when removing all assembly-based information (Fig. S3). Thus, light-induced spike patterns of peers provide more information than the single pair regarding its own STG change.

To determine whether light-induced spike patterns can predict STG changes at the assembly level, we computed an assembly CCH difference for every CA by summing all pairs in the assembly (n=122 CAs; Fig. 3K). Classifiers trained on the assembly CCH differences (Fig. 3L) yielded an AUC of 0.634 [0.610 0.658], higher than the AUC yielded by the random CAs (0.497 [0.456 0.540]; p<0.001, U-test; Fig. 3M). Thus, changes in spike timing of the entire CA predict STG changes at the assembly level.

### Precise timing carries more information than the initial conditions about changes in spike transmission

To determine temporal resolution which affects the CA STG changes, we first narrowed down the temporal precision of the predictive spike patterns by using different segments of the CCH differences for classification. The highest performance was obtained when a window of ±10 ms was centered at zero lag, yielding AUCs of 0.725 [0.706 0.751] (Fig. 4A-B). Next, we compared the contribution of co-firing (at the timescale of the ±10 ms window) and millisecond-timescale spike timing by manipulating the CCH difference vectors. To remove all co-firing (“rate”) information, we Z-scored every vector (Fig. 4C, **red**). To remove all information about precise timing without modifying co-firing information, we shuffled the order of the 1 ms bins in the CCH difference vector (Fig. 4C, **blue**). When only co-firing information was maintained, classification was at chance level (AUC, 0.506 [0.455 0.546]; p=0.85, Wilcoxon’s test; Fig. 4D, **blue**). However, when only timing information was maintained, classification yielded AUCs of 0.746 [0.718 0.763] (p<0.001; Fig. 4D, **red**). Thus, millisecond-timescale light-induced changes of spike timing within PYR-PV CAs provide information about long-term STG changes.

**Figure 4.**
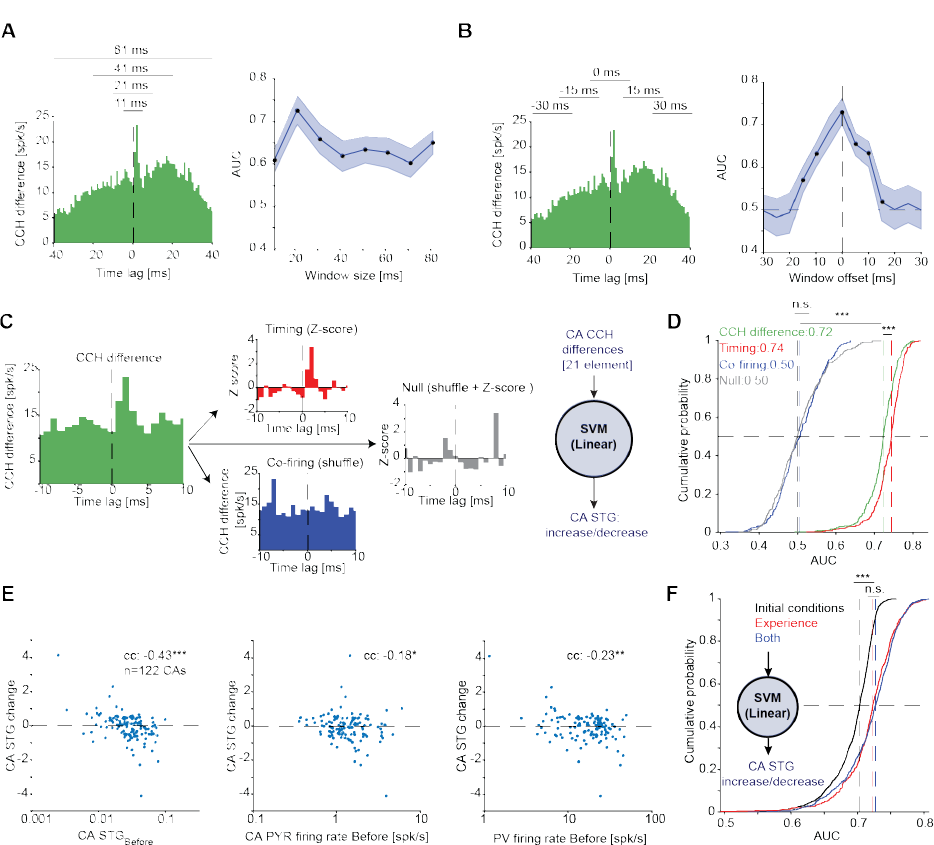
Precise timing carries more information than the initial conditions about changes in spike transmission. (**A**) **Left**. Example CA CCH difference. **Right**, Mean (solid line) and SD (band) of AUCs yielded by linear SVMs using different window sizes (n=100 repetitions). All windows are centered at zero time-lag. Here and in **B**, black points indicate p<0.05, Wilcoxon’s test corrected for multiple comparisons. (**B**) Same as **A**, for different offsets of a 21 ms sliding window. (**C**) Disambiguating the contribution of precise spike timing and co-firing to the prediction of long-term STG changes. (**D**) AUCs for four classifiers based on the same 122 CA CCH difference vectors manipulated as in **C**. Here and in **F**, n.s./***: p>0.05/p<0.001, U-test. (**E**) CAs STG change vs. three initial conditions derived from the Before epoch. */**/***: p<0.05/p<0.01/p<0.001, permutation test. (**F**) AUCs produced by three classifiers predicting CA STG increase/decrease. Black, AUCs of a classifier that received as input the CA STGBefore and the CA PYR firing rate features as in **E**. Red, AUCs yielded by a classifier that received only light-induced Experience epoch features, namely the CA CCH differences at the optimized window of ±10 ms as in **D**.

If there is a connection, the initial connectivity strength may constrain further changes^9^. Indeed, pairwise STG changes were negatively correlated with STGBefore (cc: −0.35; n=1026 PYR-PV pairs; p<0.001, permutation test; Fig. S4A). Beyond the STG itself, other initial conditions (IC) may constrain STG changes. Previous work in intact animals quantified CCH changes between PYR and interneurons as a function of firing rate changes^19^. We found that STG changes were not consistently correlated with presynaptic firing rates during the Before epoch (cc: −0.05; p=0.143; Fig. S4B), but were negatively correlated with the initial postsynaptic firing rates (cc: −0.22; p<0.001; Fig. S4C). In sum, STG changes depend on the initial conditions.

To directly assess the relative contribution of the IC and the light-induced spike patterns, we trained classifiers to predict increase/decrease of the CA STG changes. As input, we used STGBefore and PYR firing rate (Fig. 4E), which optimized the IC prediction (Fig. S4E), and the 21-element CA CCH differences, which optimized the light-induced patterns predictions (Fig. 4D-E). The classifier that used CA CCH differences alone yielded an AUC of 0.725 [0.706 0.751], higher than the classifier that used IC information alone (0.71 [0.69 0.72]; n=122 CAs; p<0.001, Wilcoxon’s test; Fig. 4F). Thus, light-induced precise spike patterns during the Experience epoch carry more information about assembly connectivity changes compared with the IC.

## Discussion

We found that precise spatiotemporal spike patterns generated by closed-loop optogenetic manipulations lead to long-lasting modifications of spike transmission among PYR-PV pairs. Spike transmission changes occur concurrently across multiple PYR-PV pairs within the same CA. Changes in STGs are more accurately predicted by spike patterns that include the same-assembly peers, compared with the specific pair. Modifications of STGs are constrained by the initial spike transmission and presynaptic firing rate, but the impact of the initial conditions is smaller than the effect of spike timing.

### Plasticity in converging assemblies

We focused on plasticity of spike transmission between PYRs and PV CAs in CA1^18, 20^. Previous research demonstrated that PYR-interneuron networks in CA1 play a crucial role in the emergence and reorganization of place fields^21, 23^ and memory regulation^24^. Furthermore, experience-dependent plasticity has been observed in neocortical^25, 26^ and hippocampal PYR-interneuron networks^26–28^. We found that STGs in the same CA are similar to begin with, i.e., “wired together”. The very organization of cells into assemblies based on postsynaptic neurons is also influenced by anatomical^29^, functional^30^, and embryological^31^ properties. Furthermore, we found that same-assembly connections undergo simultaneous same-direction modifications, i.e., “change together”. These findings are consistent with theories suggesting that neural coding is facilitated by cell assemblies linked together by dynamic changes of synaptic weights^32–36^. Our results highlight the significance of converging assemblies as natural building blocks of neuronal codes, facilitated by simultaneously changes in connectivity of CAs.

A closed-loop approach was used to pair a single spike of a presynaptic neuron with spiking activity of the postsynaptic neuron^37^. The approach differs from previous in vivo STDP experiments, which used sensory stimuli to pair the activity of a postsynaptic cell with activity of a population of presynaptic and other cells^13–15, 17, 38^. The present approach allows examining STG changes for each pair while taking into account other connections as well. The observation that STG changes occur concurrently for all converging connections may not be exclusive to PYR-PV CAs^39^. The same mechanism may underlie the plasticity reported in other in vivo STDP studies, where the spike timing of the entire CA influences all of the constituent connections.

### Timescale of plasticity

The novel short-term spike patterns between PYR-PV pairs had a long-lasting impact on the STG, highlighting the role of spike timing in synaptic plasticity. The patterns that predicted the changes occurred on a timescale of milliseconds, consistent with the effects of spike timing observed in STDP studies in vitro^9–11, 40, 41^ and in vivo^14, 15, 17, 42^. Modifying synaptic connections based on experienced spike timing at short timescales has many theoretical and practical advantages^6, 8, 43–45^.

However, highly precise patterns at short timescales leave open the question of how associations are established over behaviorally-relevant timescales spanning seconds^46^. Recent work proposed a form of plasticity that operates on a longer timescale^16, 47, 48^, “behavioral timescale synaptic plasticity” (BTSP). In BTSP, second-long plateau potentials of a single hippocampal PYR trigger the formation of place fields by altering connectivity with innervating CA3 input^16, 49^. The apparent discrepancy between the millisecond-timescale CA findings and the second-timescale BTSP results may be attributed to different intrinsic properties of PYR-PYR and PYR-PV CAs. A second manner to settle the apparent timescale discrepancy is to consider that in the intact hippocampus, long plateau potentials necessarily overlap other ongoing events. For instance, during theta oscillations, spikes of presynaptic PYR and postsynaptic INT are organized in precise sequences at millisecond timescale^50–54^. The timescale of multi-neuronal spiking during BTSP is presently unknown, since spikes of CA3 and CA1 PYRs were not recorded simultaneously. We hypothesize that BTSP requires short-timescale spiking patterns of presynaptic CA3 and postsynaptic CA1 PYR. The hypothesis may be tested by recording PYR spiking in CA3 and CA1 during the induction of CA1 place fields, potentially elucidating the timescale of spiking activity during the behaviorally-relevant alternation of synaptic connections.

## Acknowledgements

We thank György Buzsáki, Tom Manovitz, Kenji Mizuseki, Maoz Shamir, Mark Shein-Idelson, Anton Sirota, and Hadas E. Sloin for constructive comments. This work was supported by the United States-Israel Binational Science Foundation (BSF; 2015577); by the European Research Council (679253); by the Israel Science Foundation (638/16); by the Israel Science Foundation FIRST Program (1871/17); by the Rosetrees Trust (A1576); by the Canadian Institutes of Health Research (CIHR), the International Development Research Centre (IDRC), the Israel Science Foundation (ISF), and the Azrieli Foundation (2558/18); and by the Zimin Institute.

## Author contributions

L.S. and E.S. conceived and designed the study. L.S., S.So., and E.S. constructed optoelectronic probes and implanted animals. L.S., S.So., A.L., and S.Si. carried out experiments. L.S. and E.S. analyzed data. L.S. and E.S. wrote the manuscript, with input from all authors.

## Materials and Methods

### Experimental subjects

A total of seven freely-moving male mice were used in this study (Table S1). The mice aged 16 [8,30] weeks (median [range]) at the time of implantation. Animals were healthy, were not involved in previous procedures, and weighed 28.6 [22.7,30.1] g at the time of implantation. Four mice expressed ChR2 in PV cells under the PV promoter (PV::ChR2), achieved by crossing PV-Cre males (JAX #008069, The Jackson Laboratory) with ChR2 reporter females (Ai32; JAX #012569). Two mice expressed ChR2 in PYRs, generated by crossing CaMKII-Cre males (JAX #005359) with Ai32 females (CaMKII::ChR2). One mouse expressed ChR2 in Somatostatin cells, generated by a crossing SST-Cre male (JAX #013044) with an Ai32 female (SST::ChR2). Mice were single housed to prevent damage to the implanted apparatus. All animal handling procedures were in accordance with Directive 2010/63/EU of the European Parliament, complied with Israeli Animal Welfare Law (1994), and approved by Tel Aviv University Institutional Animal Care and Use Committee (IACUC #01-16-051 and #01-20-049).

### Probes and surgery

Every animal was implanted with a multi-shank silicon probe attached to a movable microdrive and coupled with optical fibers as previously described^37^. The probes used were Buzaski32 (NeuroNexus), Stark64 (Diagnostic Biochips), Dual-sided128 (Diagnostic Biochips), Dual-sided64 (Diagnostic Biochips), and Stark128 (Diagnostic Biochips). The Buzaski32 probe consists of four 52 μm wide, 15 μm thick shanks, spaced horizontally 200 μm apart, with each shank consisting of eight recording sites, spaced vertically 20 μm apart. The Stark64 probe consists of six 48 μm wide, 15 μm thick shanks, spaced horizontally 200 μm apart, with each shank consisting of 10-11 recording sites, spaced vertically 15 μm apart. The Dual-sided128 probe consists of two Stark64 probes attached back-to-back, yielding six 48 μm wide, 30 μm thick dual-sided shanks. The Dual-sided64 probe consists of two 70 μm wide, 30 μm thick dual-sided shanks, spaced horizontally 250 μm apart, with every side of each shank consisting of 16 channels spaced vertically 20 μm apart. The Stark128 probe consists of eight 33.5 μm wide, 15 μm thick shanks, spaced horizontally 125 μm apart, with each shank consisting of 16 recording sites, spaced vertically 15 μm apart. All probes were implanted in the neocortex above the right hippocampus (PA/LM, 1.6/1.1 mm; 45° angle to the midline) under isoflurane (1%) anesthesia as previously described^55^. After every recording session, the probe was translated vertically downwards by up to 70 μm. Analyses included only recordings from the CA1 pyramidal cell layer, recognized by the appearance of multiple high-amplitude units and spontaneous iso-potential ripple events.

### Recording sessions

Neuronal activity was recorded in 4.3 [3.8 4.8] hour sessions (median [IQR]). Animals were equipped with a 3-axis accelerometer (ADXL-335, Analog Devices) for monitoring head movements. Head position was tracked in real time using two head-mounted LEDs, a machine vision camera (ace 1300-1200uc, Basler), and a dedicated system^56^. Every session started with a baseline neural recording of at least 15 min, while the animal was in the home cage or in a 0.8 m diameter open field. Following the baseline recordings, a “Before” epoch that lasted a median [range] of 45 [40,45] min was carried out, during which the animals were free to behave but no light stimuli were applied. Following the Before epoch, the Experience epoch began, which lasted 55 [29,80] min. During the Experience epoch, the Stimulation group (n=31 sessions in four mice; Table S2) received closed-loop optogenetic stimuli, whereas the Control group (n=11 sessions in six mice; Table S3) underwent continuous recording without any stimuli. Following the Experience epoch, the After epoch was carried out without any stimuli, lasting 45 [40,45] min.

### Closed-loop stimulation

For the execution of closed-loop stimulation, we selected PYR spike waveform to serve as a trigger. To select the trigger PYR, we manually picked up to four same-shank channels, used for detecting and acquiring spikes over a duration of 2-3 min with a real-time digital signal processor operating at 24414 Hz (RX8, Tucker-Davis Technologies). After a batch of spikes was collected, spikes were sorted offline by applying principal component analysis (PCA) to each channel, followed by the use of the K-means algorithm on the three first PCA components extracted from each channel to form clusters. We then selected a spike cluster for parameter extraction during real-time detection. We manually selected three voltage windows, using the same channels used for real-time spike detection. Each window consisted of two voltage points and one time point. After uploading the definitions to the digital signal processor, only spikes that passed through all the windows were detected. Upon the real-time detection of a PYR spike, a voltage command was issued to a linear current source, prompting the delivery of a 30 ms light stimulus. Post-hoc analysis indicated that light stimuli were given a median [IQR] of 3 ms [3 3] ms (n=394,886 light stimuli) after the spike trough occurred. Following a light stimulus, a dead-time of 20 ms was applied during which no stimulation was given.

### Spike detection, spike sorting, and cell type classification

Neural activity was filtered, amplified, multiplexed, digitized on the headstage (0.1–7500 Hz, x192; 16 bits, 20 kHz; RHD2132 or RHD2164, Intan Technologies), and recorded by an RHD2000 evaluation board (Intan Technologies). Offline, spikes were detected and sorted into single units automatically using either KlustaKwik3^57^ or KiloSort2^58^. Automatic spike sorting was followed by manual adjustment of the clusters. Only well-isolated units were used for further analyses (amplitude >40 μV; L-ratio <0.05; inter-spike interval index <0.2; ^20^). Units were then classified into putative PYR or PV-like interneurons (INT) using a Gaussian mixture model^59^.

### Selection of a subset of data

To select PYR-PV pairs for the Stimulation and the Control groups, we applied the following criteria: (1) Detection of an excitatory monosynaptic connection in the CCH using spike trains from the combined Before and After epochs (p<0.001, Poisson test). (2) Accumulation of at least 400 counts in the count CCHs of each of the Before and After epochs at the −30<τ≤ 30 range. (3) Identification of the excitatory monosynaptic peak (0<τ≤ 5 range) as the highest peak in the Before or After CCH. The Stimulation group included only pairs in which the postsynaptic cell was a PV cell (i.e., activated by the closed-loop stimulation).

The Stimulation group consisted of 1838 PYRs and 420 INT recorded during 31 sessions from four mice, forming 24461 pairs. Of these, 3702 (15%) PYR-INT pairs were connected (p<0.001, Poisson test). After applying the above criteria, the group consisted of 689 PYRs, 122 PV cells, and 1026 connected pairs from 29 sessions in the four mice. The Control group consisted of 547 PYRs and 152 INT recorded during 11 sessions from six mice, forming 10743 pairs. Of these, 1197 (11%) pairs were connected. After applying the criteria, the group consisted of 224 PYRs, 48 INT, and 388 connected pairs from 9 sessions in five mice.

### Quantification of closed-loop feedback: closed-loop efficiency and light response

To determine what fraction of spikes of a given unit were used to generate light stimuli, we defined a “closed-loop efficiency” (CLE) measure. To estimate the CLE, we used the same approach as for computing the STG (see below), based on the peristimulus time histograms (PSTHs, 1 ms bin size) instead of the CCH. The PSTHs were constructed around stimulus onset for every PYR in the Stimulation group and scaled to spk/s (as in Fig. 3A). The CLE was the calculated as the area under the peak in the −5≤τ<0 ms ROI, and the baseline activity was determined by hollowed median filtering (5 ms halfwidth) of the PSTH. The CLE is limited to the [0,1] range. A CLE of zero indicates that no spikes were followed by light stimuli, and a CLE of 1 indicates that every spike was followed by a single light stimulus. In practice, the maximal CLE is lower than 1 even for perfect detection due to the processing delay (3 ms), stimulus duration (30 ms), and post-stimulus dead-time (20 ms). The CLEs were 0.004 [0,0.848] (median [range]; n=689 PYRs). In trigger PYRs, defined as PYRs with consistent peak in the −5≤τ<0 ms ROI of the PSTH (p<0.001, Poisson test), the CLEs were 0.054 [0.002,0.848] (n=198; Fig. S2C).

To identify PV cells that were light-activated by the closed-loop stimulation, we constructed PSTHs (bin size of 1 ms) around the stimulus onset for every PV cell (as in Fig. 1J). PV cells that exhibited a consistent increase in firing rate during the 10≤τ<30 ms interval relative to baseline activity (p<0.05, Poisson test), were classified as light-activated PV cells. The baseline activity was determined by the mean firing rate in the 15 ms preceding light onset.

### Computing STG changes

To compute STG changes, two count cross-correlation histograms (CCHs; 0.5 ms bins) were constructed for every pair, separately for the Before and for After epochs. The STG was then computed for each of the two CCHs as previously described^22^. Briefly, the spike transmission curve was estimated by the difference between the deconvolved CCH and the baseline, determined by hollowed median filtering of the count CCH, scaled to spk/s. The STG was defined as the area under the peak in the monosynaptic temporal region of interest (ROI; 0<1≤5 ms), extended until the causal zero-crossing points. The STG change was then computed as the base-2 logarithm of the ratio between the STGAfter and the STGBefore. The STG change is not limited to a specific range. An STG change of 1 (or −1) indicates that the STG has doubled (or halved) following the “Experience” epoch, while a value of 0 indicates no change in STG.

To assess the consistency of changes in STG (i.e., a significant increase or decrease), we conducted a permutation test. First, we generated a binary spike time lag matrix (0.5 ms bins) for each PYR-PV pair using spikes that occurred during the Before and After epochs. Each row in the matrix represented a single PYR spike, and every column denoted a time lag. The value in every cell indicated whether a PV spike occurred (1) or not (0) during the corresponding time interval. Every row was labelled according to the source spike (Before or After). The original STG change was computed from the count CCHs generated by summing the rows that correspond to the Before epoch and the After epoch separately. Second, we shuffled the row labels of the matrix and partitioned the matrix into two matrices with identical sizes to the original matrices, but according to the shuffled labels. We then computed the STG change using the two shuffled matrices. Third, we repeated the shuffling process 2000 times and obtained a distribution of STG changes, which was used to estimate the two-tailed p-value of the original (observed) STG change. Specifically, the p-value was the fraction of shuffled STG changes that were more extreme than the observed STG change (i.e., either higher or lower). The fraction was calculated by adding 1 to the number of more extreme STG changes and dividing by the total number of shuffles plus 1.

We found that 11% (116/1026) of the Stimulation pairs exhibited a consistent STG increase (p<0.001, Binomial test comparing to chance level, 2.5%), and 16% (160/1026) exhibited a decrease (p<0.001). Similar results were also observed in the Control pairs group, recorded during long no-stimulus durations. Of the control pairs, 14% (54/388) exhibited a consistent STG increase (p<0.001), and 12% (45/388) exhibited a decrease (p<0.001).

### Synchrony quantification

To quantify synchrony between a pair of PYR spike trains s1 and s2, we constructed the count CCH using 0.5 ms bins. We then counted the number of coincident counts in the synchrony ROI (−1::;1::;1 ms) nsync, and divided that by the geometric mean total number of spikes in each of the two trains, 17sync = nsync/N1N2. The 17sync measure generalizes the synchrony effect size of^60^ to the symmetric setting. 17sync is bounded 0::;17sync::;1, equaling 1 when all spikes of one of the trains are synchronous with the other train. Notably, 17sync may be non-zero simply by chance, even when using a small bin size and a small ROI, and even when there is no millisecond-timescale synchrony. To obtain an estimate of the synchrony above expected by chance, we defined chance level using timescale separation, by hollowed median filtering (5 ms halfwidth) of the CCH, obtaining a predictor CCH, pred^22^. We then derived the chance level synchrony effect size from the predictor in the same manner, 17pred = npred/✓N1N2. The synchrony measure, 1¢17 = 17sync-17pred, is bounded −1::;1¢17::;1. To determine significance, we used a Poisson test, estimating the Poisson probability of observing nsync or more counts when npred are expected and applying a continuity correction^60^.

We measured pairwise synchrony for every PYR in the set of n=689 PYRs during the entire recording session (excluding stimulation times) using the abovementioned bin size and ROI. After excluding same-shank PYRs, the mean synchrony was computed for the target PYR and all peers (PYRs that participated in the same CA) and for all non-peer PYRs. We also counted the fraction of synchronized peers and synchronized non-peers (p<0.001, Poisson test). A total of 623/689 PYRs had at least one peer and one non-peer recorded on other shanks. The median [IQR] number of peers per PYR was 10 [6 19], and the median [IQR] mean synchrony with peers 0.0014 [0.0008 0.0025] (n=623 PYRs). For non-peers there were 16 [8 29] non-peers per PYR, and the mean synchrony was lower at 0.0010 [0.0007 0.0016] (p<0.001, Wilcoxon’s paired test; Fig. S1I). The fraction of synchronized peers per PYR was 26% [8% 46%], higher than the fraction of synchronized non-peer pairs per PYR (16% [55% 36%]; (Wilcoxon’s paired test; Fig. S1J). Considering all pairs, the fraction of synchronized peer pairs was 31% (2398/7692 pairs), higher than the overall fraction of synchronized non-peer pairs (2239/11211 pairs, 20%; p<0.001, G-test;). These results were not sensitive to specific parameter values, and similar (and significant) results were obtained for bin sizes of {0.25, 0.5, 1} ms, and for ROIs of {-0.5::;1::;0.5, −1::;1::;1, −1.5::;1::;1.5} ms.

### Cross-validated regression

To predict a single STG change (Fig. 2M-O), we utilized a support vector regressor (SVR) with a radial basis function kernel. The inputs for the SVR were different combinations of the following three features: (1) the mean STG changes of peer pairs, (2) the mean STG changes in non-peer pairs, and (3) changes in the firing rate of the postsynaptic cell. We conducted training and testing of the SVR using a five-fold cross-validation method. The entire process was reiterated 100 times, each instance partitioning the data into five random segments. The coefficient of determination (R^2^) for each 5-fold cross-validation was computed to evaluate the performance under cross-validation. The control SVR used as an input the mean STGs of pairs from the same session, which were randomly shuffled into peer pairs and non-peer pairs groups (Fig. 2M-N). To assess the individual contribution of peers, non-peers, and the firing rate changes for predicting the single STG change, we repeated the process by training additional SVRs, each with only two of the features (Fig. 2O).

### Binary classification

To predict whether the STG change increased or decreased following the Experience (i.e., between the Before and After epochs), we employed a linear support vector machine (SVM). To evaluate classification performance, we trained every classifier using five-fold cross-validation, repeating the process 100 times with the data split into random five folds each time. We then calculated the AUC of each five-fold cross-validation. AUC values were compared to the corresponding values yielded by a control SVM. The control SVM used shuffled labels (Fig. 3J, Fig. S3, and Fig. S4D), shuffled CAs (Fig. 3M), or Z-scored CCH difference with shuffled bins (Fig. 4D).

Because STGs in a given CA change together (Fig. 2), the input used for the SVM classifier may contain information about the association of individual pairs to CAs, which could impact the prediction. To eliminate any assembly-based information from the classification process, we used two approaches: entire CA cross-validation or subsampling. In the first approach, entire CA cross-validation (Fig. S3A), we divided the data into five folds of identical sizes, subject to the constraints of the data, with each fold containing data from the entire CAs. In other words, pairs from the same assembly were used either for training or for testing, but never for both. Thus, the training and testing were applied to data from different assemblies. In the second, subsampling approach (Fig. S3B), we randomly selected one pair from each CA, resulting in 116 pairs. We then trained an SVM using many different partitions of random five-fold cross-validation with these 116 pairs. We repeated the process until every pair was sampled at least 100 times. As some pairs were sampled more than others, we randomly selected 100 predictions for each pair.

### Statistical analyses

In all statistical tests, a significance threshold of α=0.05 was used. An exception was the threshold used for determining whether two units exhibit monosynaptic connectivity or synchrony (α=0.001). In all cases, non-parametric testing was used. All statistical details (n, median, IQR, range, mean, SD) can be found in the main text, figures, figure legends, and tables. To estimate whether fractions were larger or smaller than expected by chance, an exact Binomial test was used (two-tailed). Differences in the proportions of two categorical variables were tested with a likelihood ratio (G-test). Differences between two group medians were tested with either Mann-Whitney’s U-test (unpaired samples) or Wilcoxon’s paired signed-rank test (two-tailed). To estimate whether a median was larger or smaller than expected by chance, a Wilcoxon’s signed-rank test was used (two-tailed). Association between parameters was quantified using Spearman’s rank correlation and tested with a permutation test. For all figures, n.s.: p> 0.05; *: p<0.05; **: p<0.01; ***: p<0.001.

---

**Figure S1.**
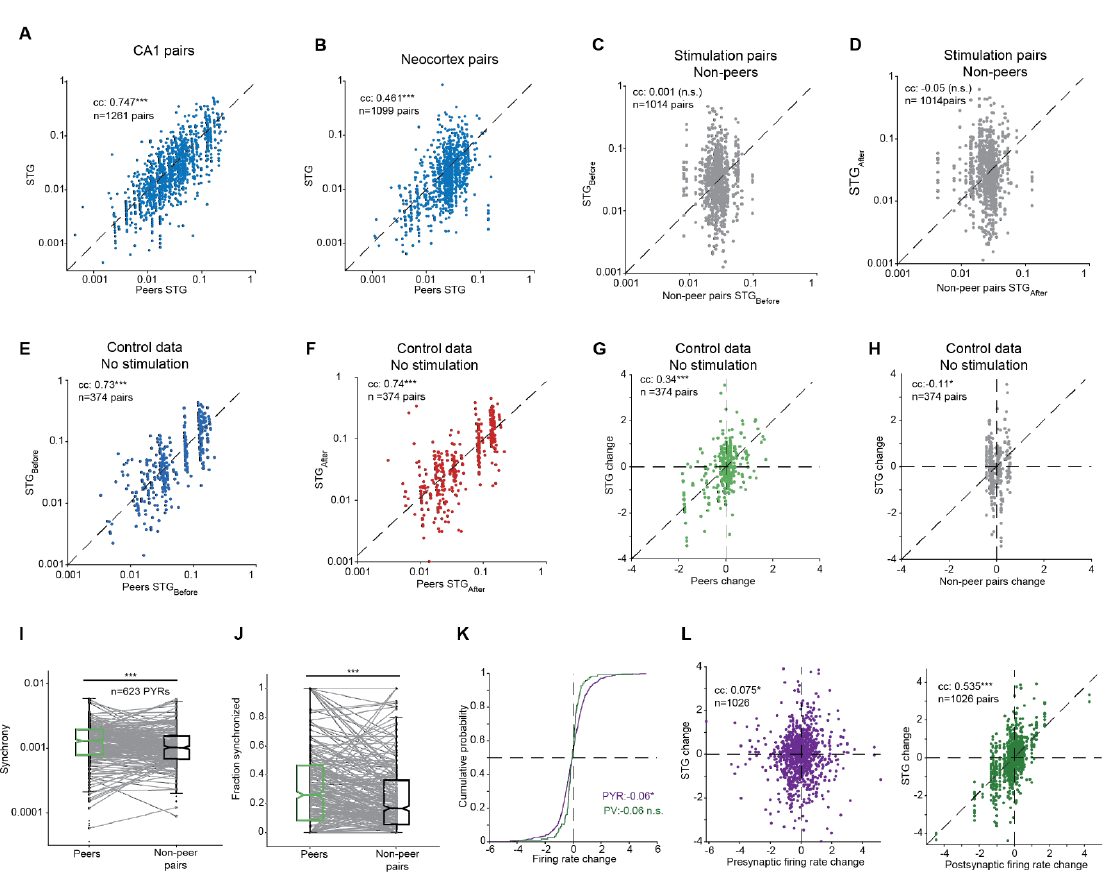
STGs are similar for pairs in the same converging assembly. (**A**) STG of every CA1 PYR-INT pair vs. mean STG of all peers. Peers, PYR-INT pairs that share the same postsynaptic INT, forming a CA. n=1261 pairs in 139 CAs recorded from hippocampal region CA1 of three mice^22^. Here and in **B**-**H**, and **K**, n.s./*/**/***: p>0.05/p<0.05/p<0.01/p<0.001, permutation test. (**B**) STGs of neocortical PYR-INT pairs vs. mean peers STG. n=1099 pairs in 100 CAs recorded from six mice^20^. (**C**) The STGBefore of every pair vs. the mean STGBefore of all non-peer pairs. Here and in **D**, data are shown for the stimulated 1014/1026 PYR-PV pairs as in Fig. 2C. (**D**) Same as **C** for the STGAfter. (**E**) The STGBefore of every pair vs. the mean STGBefore of all peers. Here and in **F,G,H**, data are shown for the 374/388 PYR-PV Control pairs which did not undergo light stimulation during the “Experience” epoch. (**F**) Same as **E** for the STGAfter. (**G**) STG change vs. the mean STG change of all peers. (**H**) Same as **G** for non-peer pairs. (**I**) Mean synchronous firing between a single PYR and either other peers PYRs or other non-peers PYRs. n=623/689 PYRs which have both peers and non-peers PYR recorded on a different shank. Here and in **J**, ***: p<0.001, Wilcoxon’s paired test. (**J**) Same as **I** for the fraction of synchronized peers and synchronized non-peers of each PYR. (**K**) Distribution of the firing rate change for PYRs (n=689) and PV cells (n=122). The firing rate change is defined as the base-2 logarithm of the ratio between the firing rate during the After epoch and the firing rate during the Before epoch. Median [IQR] values are −0.06 [−0.61 0.47] for the PYRs, and 0.06 [−0.32 0.18] for the PV cells. n.s./*: p>0.05/p<0.05, Wilcoxon’s test comparing to a zero-change null. (**L**) STG changes vs. firing rate changes of the presynaptic or the postsynaptic cell.

**Figure S2.**
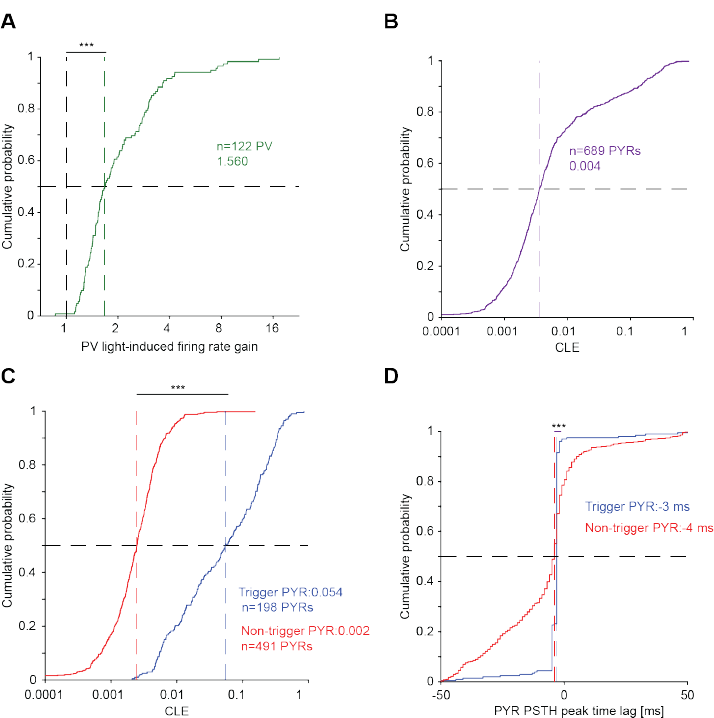
PYR spikes trigger closed-loop light stimulation with millisecond timescale resolution. (**A**) Distribution of the light-induced firing rate gain for the n=122 PV cells. The light-induced gain is defined as the ratio between the mean firing rate during light stimuli and the mean firing rate during the 15 ms before stimulation onset. ***: p<0.001, Wilcoxon’s test comparing to a unity gain null. (**B**) Distribution of closed-loop efficiency (CLE) for all PYRs. For a given light-triggered PSTH, the CLE is defined as the area under the maximal peak, above baseline firing activity. (**C**) PYRs which exhibit a consistent PSTH peak within the closed-loop ROI (−5≤τ<0 ms before stimulation onset; p<0.001 Poisson test) are categorized as “trigger” cells. Here and in **D**, ***: p<0.001, U-test. (**D**) Distribution of PSTH peak time lag for trigger (n=198) and non-trigger PYRs (n=491). SDs are 8 ms for trigger PYRs and larger (14 ms) for non-trigger PYRs (p<0.001, permutation test).

**Figure S3.**
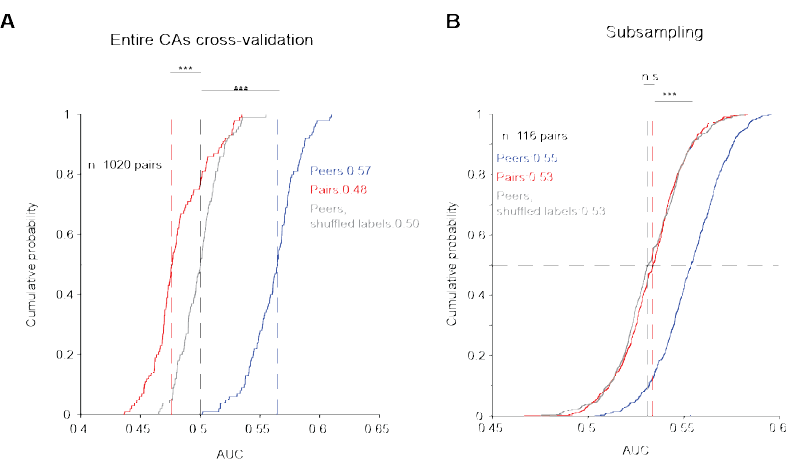
Pairwise STG changes can be predicted more accurately by peer pairs spike timing even when assembly structure information is removed. (**A**) For removing all assembly-based information from the classification process, we utilized two approaches. In the first approach, we trained cross-validated classifiers with carefully-constructed cross-validation folds. Specifically, each fold in the training and test sets utilized only entire assemblies, ensuring that pairs belonging to the same assembly were never used in both the training or test sets. We trained three types of five-fold cross-validated linear SVMs: pairs, peer pairs, and peers with shuffled labels. The process was repeated 100 times, and every iteration employed random partitions which were composed only from entire assemblies. Here and in **B**, n.s./***: p>0.05/p<0.001, U-test. (**B**) As an alternative approach for removing all assembly-based information from the classification process, we sampled only one pair from every CA. We trained three five-fold cross-validated SVMs: pairs, peer pairs, and peers with shuffled labels. Every classifier was trained during 100 independent runs, and in every run a random subset of 116/1020 pairs, one from every CAs, was employed.

**Figure S4.**
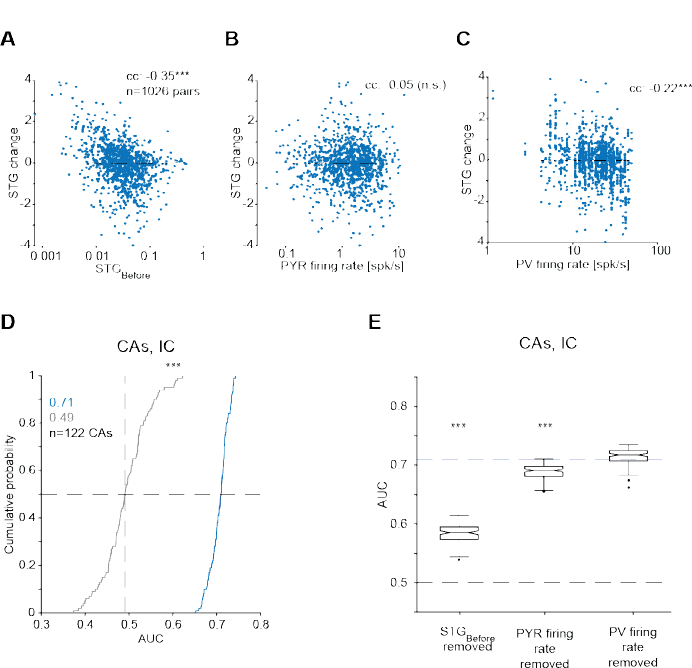
Initial conditions carry information about CA STG changes. (**A-C**) Pairwise STG change vs. three initial conditions derived from the Before epoch. n.s./***: p>0.05/p<0.001, permutation test. (**A**) STGBefore. (**B**) The mean firing rate of the presynaptic PYR. (**C**) The mean firing rate of the postsynaptic PV cell. (**D**) AUCs produced by a cross-validated binary classifier (linear SVM) trained with all three mean CA initial conditions features shown in Fig. 4E. Here and in **E**, ***: p<0.001, Wilcoxon’s test. (**E**) AUCs for three classifiers. For every classifier, a different feature was removed. Top dashed line, median AUC of the full model.

**Table S1.**
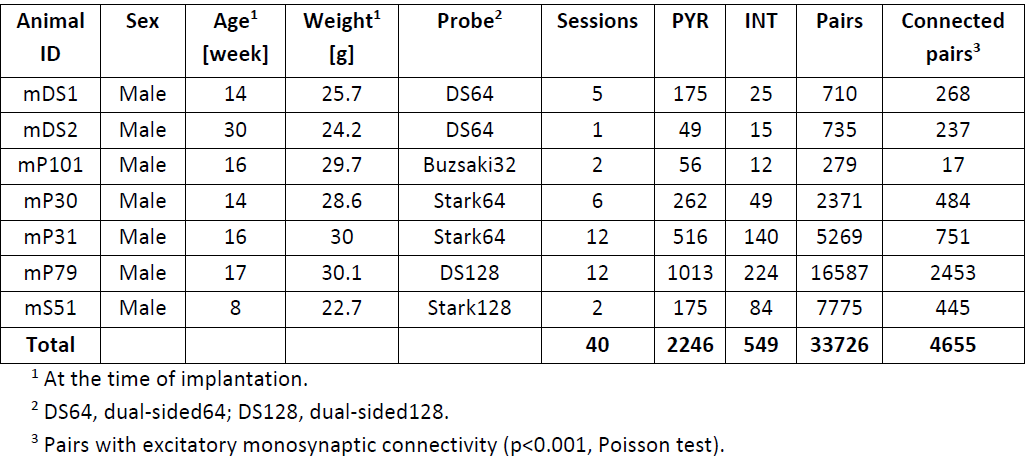
List of experimental animals.

| **Animal ID** | **Sex** | **Age<sup>1</sup>**<br>**[week]** | **Weight<sup>1</sup>**<br>**[g]** | **Probe<sup>2</sup>** | **Sessions** | **PYR** | **INT** | **Pairs** | **Connected pairs<sup>3</sup>** |
| --- | --- | --- | --- | --- | --- | --- | --- | --- | --- |
| mDS1 | Male | 14 | 25.7 | DS64 | 5 | 175 | 25 | 710 | 268 |
| mDS2 | Male | 30 | 24.2 | DS64 | 1 | 49 | 15 | 735 | 237 |
| mP101 | Male | 16 | 29.7 | Buzsaki32 | 2 | 56 | 12 | 279 | 17 |
| mP30 | Male | 14 | 28.6 | Stark64 | 6 | 262 | 49 | 2371 | 484 |
| mP31 | Male | 16 | 30 | Stark64 | 12 | 516 | 140 | 5269 | 751 |
| mP79 | Male | 17 | 30.1 | DS128 | 12 | 1013 | 224 | 16587 | 2453 |
| mS51 | Male | 8 | 22.7 | Stark128 | 2 | 175 | 84 | 7775 | 445 |
| **Total** |  |  |  |  | **40** | **2246** | **549** | **33726** | **4655** |

<sup>1</sup> At the time of implantation.
<sup>2</sup> DS64, dual-sided64; DS128, dual-sided128.
<sup>3</sup> Pairs with excitatory monosynaptic connectivity ( $p < 0.001$ , Poisson test).

**Table S2.** Units and pairs used in the Stimulation analyses.

| **Animal ID** | **All sessions** | **Valid<sup>1</sup> sessions** | **PYR** | **PV<sup>2</sup>** | **Pairs** | **STG increase<sup>3</sup>** | **STG decrease<sup>3</sup>** | **CAs** |
| --- | --- | --- | --- | --- | --- | --- | --- | --- |
| mP101 | 1 | 1 | 9 | 3 | 10 | 4 | 2 | 2 |
| mP30 | 6 | 6 | 98 | 20 | 142 | 12 | 19 | 19 |
| mP31 | 12 | 10 | 163 | 33 | 210 | 19 | 21 | 31 |
| mP79 | 12 | 12 | 419 | 66 | 664 | 81 | 118 | 64 |
| **Total** | **31** | **29** | **689** | **122** | **1026** | **116** | **160** | **116** |

<sup>1</sup> Sessions with at least one valid connected PYR-PV pair.
<sup>2</sup> Units with firing rate increase during closed-loop illumination ( $p < 0.05$ , Poisson test).
<sup>3</sup> Pairs with consistent STG increase/decrease ( $p < 0.025$ , permutation test).

**Table S3.** Units and pairs used in the Control analyses (Fig. 1M)

| **Animal ID** | **All sessions** | **Valid<sup>1</sup> sessions** | **PYR** | **PV<sup>2</sup>** | **Pairs** | **STG increase<sup>3</sup>** | **STG decrease<sup>3</sup>** | **CAs** |
| --- | --- | --- | --- | --- | --- | --- | --- | --- |
| mDS1 | 5 | 4 | 52 | 9 | 83 | 5 | 8 | 6 |
| mDS2 | 1 | 1 | 41 | 5 | 93 | 22 | 15 | 5 |
| mP101 | 1 | 0 | 0 | 0 | 0 | 0 | 0 | 0 |
| mP31 | 1 | 1 | 20 | 5 | 29 | 4 | 3 | 4 |
| mP79 | 1 | 1 | 46 | 9 | 80 | 8 | 8 | 8 |
| mS51 | 2 | 2 | 65 | 20 | 103 | 15 | 11 | 17 |
| **Total** | **11** | **9** | **224** | **48** | **388** | **54** | **45** | **40** |

<sup>1</sup> Sessions with at least one valid connected PYR-INT pair.
<sup>2</sup> Units with firing rate increase during closed-loop illumination ( $p < 0.05$ , Poisson test).
<sup>3</sup> Pairs with consistent STG increase/decrease ( $p < 0.025$ , permutation test).

